# The Implications of Small Stem Cell Niche Sizes and the Distribution of Fitness Effects of New Mutations in Aging and Tumorigenesis

**DOI:** 10.1101/032813

**Authors:** Vincent L. Cannataro, Scott A. McKinley, Colette M. St. Mary

## Abstract

Somatic tissue evolves over a vertebrate’s lifetime due to the accumulation of mutations in stem cell populations. Mutations may alter cellular fitness and contribute to tumorigenesis or aging. The distribution of mutational effects within somatic cells is not known. Given the unique regulatory regime of somatic cell division we hypothesize that mutational effects in somatic tissue fall into a different framework than whole organisms; one in which there are more mutations of large effect. Through simulation analysis we investigate the fit of tumor incidence curves generated using exponential and power law Distributions of Fitness Effects (DFE) to known tumorigenesis incidence. Modeling considerations include the architecture of stem cell populations, i.e., a large number of very small populations, and mutations that do and do not fix neutrally in the stem cell niche. We find that the typically quantified DFE in whole organisms is sufficient to explain tumorigenesis incidence. Further, due to the effects of small stem cell population sizes, i.e., strong genetic drift, deleterious mutations are predicted to accumulate, resulting in reduced tissue maintenance. Thus, despite there being a large number of stem cells throughout the intestine, its compartmental architecture leads to significant aging, a prime example of Muller’s Ratchet.

## Introduction

### Evolution in Somatic Tissue

The epithelial tissues within many animals are continually replenished by populations of stem cells that divide throughout the organism’s lifetime. For instance, the epithelial lining of the intestinal tract is replaced weekly by millions of independent populations of stem cells located in intestinal crypts (reviewed in Barker (2014)). This continual division provides an opportunity for mutation, resulting in the accumulation of mutant lineages and somatic evolution (Lynch, 2010). Stem cell lineages with decreased fitness, or a diminished ability to divide and survive, will represent a failure in this tissue renewal process and the aging of tissues and multicellular organisms as a whole (López-Otín et al., 2013; Moskalev et al., 2013). Lineages with increased fitness, or faster division rates and an increased propensity to survive, will result in the accumulation of cells and neoplasia (Merlo et al., 2006). Although considered premalignant at the onset, the accumulation of cells into a polyp, in which cells continually divide and accumulate subsequent mutations, can develop a cancerous phenotype over time (Winawer, 1999).

### Distribution of Fitness Effects

The effect that a new mutation will have on an individual’s fitness can be characterized by a distribution of fitness effects (DFE). The DFE of several organisms have been experimentally estimated using mutation accumulation experiments or directed mutagenesis experiments in the laboratory (Eyre-Walker and Keightley, 2007; Halligan and Keightley, 2009). The majority of random mutations to a genome that affect fitness have a deleterious effect on fitness, while a small subset increase fitness (Eyre-Walker and Keightley, 2007). Additionally, many mutations that affect fitness have a small effect, while few have a large effect. In general, both beneficial (Imhof and Schlotterer, 2001; Kassen and Bataillon, 2006; Orr, 2003) and deleterious (Elena et al., 1998) mutational fitness effects can be described well using an exponential distribution, although there are exceptions (Rokyta et al., 2008) and compound distributions or distributions with more parameters may better fit empirical measures of DFE (Sanjuán et al., 2004).

By understanding the mutational DFE in somatic tissue we can predict the evolutionary trajectories of tissues within multicellular organisms as they age. In this work, we consider mutations to a stem cell’s division rate and the rate at which it commits to differentiation. Fitness in this context refers to the eventual size of the stem cell’s lineage throughout the crypt. Increases in division rate and decreases in differentiation rate are therefore beneficial. Due to the specific structure of the stem cell population, we note that increased fitness does not necessarily imply an increased likelihood that a given beneficial mutation will fix in the stem cell pool. Indeed, beneficial mutations in division rate lead to an increased fixing probability, while beneficial mutations in differentiation rate are neutral in this sense. Although there has been no direct measurement of the distribution of fitness effects in somatic tissue (but see Vermeulen et al. (2013); Snippert et al. (2014) for estimations of the selective advantage for some known cancer drivers), the evolution of cancer progression has been previously modeled using discrete (Beerenwinkel et al., 2007; Bozic et al., 2010; McFarland et al., 2013) and continuous (Foo et al., 2011) fitness effects. Here, we differ from these previous models by investigating different mutational effect frameworks using parameters derived from whole organisms to explore mutation accumulation in crypts initialized at their measured healthy size in mice and humans and quantify both aging and tumorigenesis.

When quantifying tumor incidence we are concerned with the moment that the regulatory regime in the intestinal crypt breaks down: when the stem cell division rate exceeds its differentiation rate. We call this point the *tumorigenesis threshold*. The resulting population will accumulate stem cells without bound, which is thought to be the cause of crypt fission and the main mechanism of polyp or adenoma growth (Wong et al., 2002; Loeffler and Grossmann, 1991). We investigate the full spectrum of deleterious and beneficial mutational effects on the progression of a healthy crypt to tumor initiation using empirically measured rates of division.

The evolution of multicellularity has necessitated the evolution of regulatory systems that hold somatic stem cells at a relatively low growth rate, i.e. fitness, (when compared to their maximum potential) in order to ensure the cooperation of the different cellular systems constituting a whole organism. As such, in addition to the beneficial mutations that would be accounted for by a typical DFE for whole organisms (which are commonly assumed to already be highly fit (Orr, 2010)), we expect mutations of large effect in somatic tissue as regulatory processes become dysfunctional, such as the deactivation of tumor suppressor genes or the activation of oncogenes. It is reasonable to hypothesize that a heavy-tailed distribution could better classify mutational effects that have a beneficial effect in somatic stem cells by capturing both the mutations of small effect and also having a non-trivial probability of capturing the mutations of large effect often associated with cancer.

We evaluate whether or not the DFE estimated in whole organisms can explain known tumor incidence in the intestine. Further, we explore whether or not tumor incidence is better explained by a heavy-tailed distribution for mutations beneficial to fitness. Thus, we create a model of an evolving intestinal stem cell pool and implement alternate DFE and compare the resultant incidence curves to known tumor incidence curves.

## Materials and Methods

### Description of the Model

**Crypt Population Structure**. The base of each intestinal crypt harbors a population of symmetrically dividing cells expressing markers associated with the stem cell phenotype (Lopez-Garcia et al., 2010; Snippert et al., 2010). Within this population there exists a subpopulation niche that is responsible for maintaining tissue homeostasis (Kozar et al., 2013; Vermeulen et al., 2013). We model the stem cells of the intestinal crypt as two populations of cells, the first being this stem cell niche, which consists of a fixed population of stem cells, *N*, and the second consisting of the stem cells displaced from this niche but not yet committed to differentiation. The sum of these populations represent the total number of stem cells within the crypt, *N_T_*. In our model, cells within the niche divide at rate *λ* and displace their neighbors through overcrowding, as proposed by Lopez-Garcia et al. (2010) and revealed by live *in vivo* imaging by Ritsma et al. (2014). This population of cells experiences genetic drift and selection; cells that have a higher division rate are more likely to push their neighbors out of the niche (as demonstrated by Snippert et al. (2014)) and cells with lower division rates are more likely to be displaced. Mutations may occur at division with mutation rate *μ* and result in either a lineage with a new division rate or a lineage with a new rate of committing to differentiation. Displaced stem cells divide at the rate of their progenitor cells in the niche and commit to differentiation at rate *ν*, hereafter referred to as the differentiation rate. We assume that once a lineage commits to differentiation it is destined to be expelled from the crypt. We define tumorigenesis in the crypt as the moment a lineage of stem cells with a division rate greater than its differentiation rate has become fixed in the niche, resulting in exponential population growth. We note that, although a stem cell’s propensity to commit to differentiation in healthy tissue is partially dependent on external signaling queues, such as Wnt signals from Paneth cells in the small intestinal crypt stem cell niche (Clevers, 2013), the ability of a stem cell to interpret and respond to, or even gain independence from, external signals is an intrinsic and heritable property of the stem cell (Reya and Clevers, 2005). The parameters *λ*, *N*, and *N_T_* have been previously estimated (Kozar et al., 2013; Vermeulen et al., 2013), and we calculate *ν* according to equation 4 in the Appendix: Description of the mathematical methodology, where 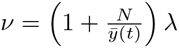 and 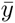 is the average number of stem cells outside of the niche.

**Distribution of Fitness Effects**. We first describe our model of mutations that affect the division rate of stem cells and address mutations that affect differentiation rate later in section “Mutations that alter the differentiation rate of stem cells result in rapid aging and tumorigenesis” When mutations occur the new division rate is greater than the previous rate with probability *P_B_*, and the mean positive change of rate is *s*_+_. We consider positive and negative changes that are exponentially distributed for deleterious effects and exponentially or Pareto distributed for beneficial effects, see the Appendix: Description of the mathematical methodology. The mean negative change is *s_−_*. We define the exponential DFE in Equation 1 and the power-law DFE in Equation 2.

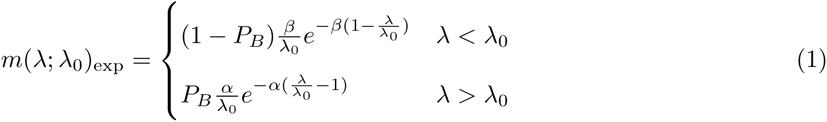

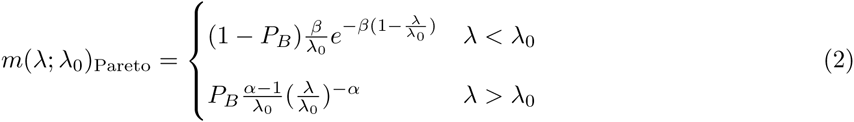

The power law distribution is well defined if *α >* 1 and is considered to be heavy-tailed (having infinite variance) if 1 < *α <* 3.

**Selection Assumptions**. We are concerned with the mutations that arise and reach fixation within the stem cell niche. Due to drift, all stem cells with the same division rate as the background population have an equal probability of reaching fixation, commonly referred to as neutral drift dynamics (Lopez-Garcia et al., 2010; Snippert et al., 2010). Following Wodarz and Komarova (2005) we use a Moran model to estimate the probability that a mutant lineage fixes in the stem cell niche:

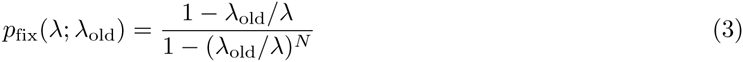
where *N* is the number of cells in the niche. The mutation rate is low relative to the division rate, so we assume that there are at most two competing division rates at any given time.

Using the above formula (3), we can use Bayes’ Theorem to compute the probability density Φ(λ | λ_old_) of a new fixed division rate λ given that the previous division rate is λ_old_:

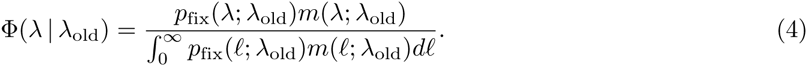

As described above, tumorigenesis occurs when the division rate λ is greater than differentiation rate *ν* and we define the point at which this happens to be the tumorigenesis threshold. In our modeling framework, each new fixed mutation presents a new possibility that the division rate exceeds the threshold for tumorigenesis. From (4) we can iteratively derive the sequence of functions {*f_n_*} that represent the density of the distribution of the stem cell division rates conditioned that *n* mutations have fixed in the stem cell niche and tumorigenesis has not occurred as of mutation *n* − 1. If we let λ_0_ denote the initial stem cell division rate, then *f*_1_(λ) = Φ(λ | λ_old_ = λ_0_). For each *n* let *p_n_* denote the probability that tumorigenesis occurs due to the nth mutation (given that *n* mutations have occurred). Then 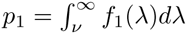. From there, we can write the recursive formulae

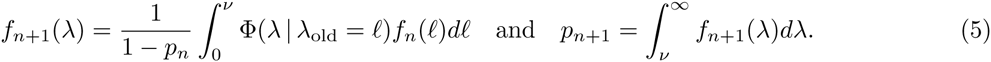

From this, we have a recursive formula for the probabilities, {*q_n_*}, that tumorigenesis has *not* occurred given *n* fixed mutations: *q*_1_ = 1 − *p*_1_ and

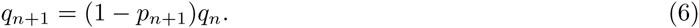

To translate this result to an individual’s lifetime, we model the time-dependent arrival of new mutations as a Poisson process with fixed rate parameter *μ* mutations per cell division. We keep track of the time-dependent number *M*(*t*) of mutations that *fix* in the stem cell niches by time *t*. Then, using *T* to denote the time that tumorigenesis occurs in a given crypt, we can write the probability that tumorigenesis has *not* occurred as of time *t* by the equation

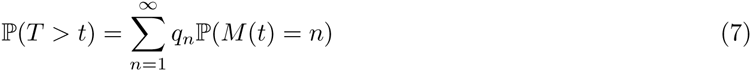

Depending on the species, an individual has hundreds of thousands or even millions of crypts. The probability that an individual has at least one crypt that has undergone tumorigenesis can be calculated by considering the distribution of fixed mutations that have accumulated among the individual’s crypts and the probability that these mutations result in tumorigenesis. This can then be extrapolated to the incidence rate of tumors among a population of individuals (Fig. 1). Let 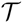 represent the time that tumorigenesis first occurs in any of an individual’s crypts. We use the following estimate to calculate tumorigenesis incidence data reported in the Results section. In the Supporting Information we describe the full calculation and a few simplifying assumptions we make to develop a computationally tractable model.

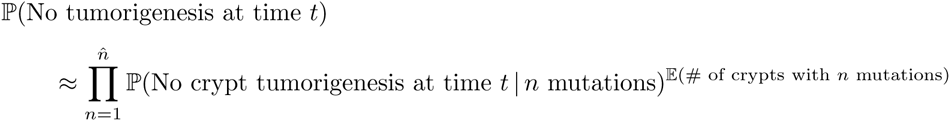

i.e.

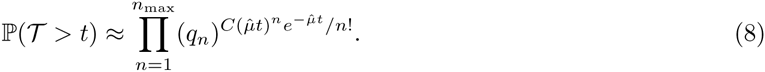

In the above, *C* is the number of crypts in the length of intestine being investigated, *n_max_* is the maximum number of mutations simulated and 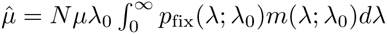, where, as above, *N* is the number of cells in the stem cell niche, *μ* is the mutation rate per cell division and *p*_fix_ is defined by Eq. 3.

**Figure 1.**
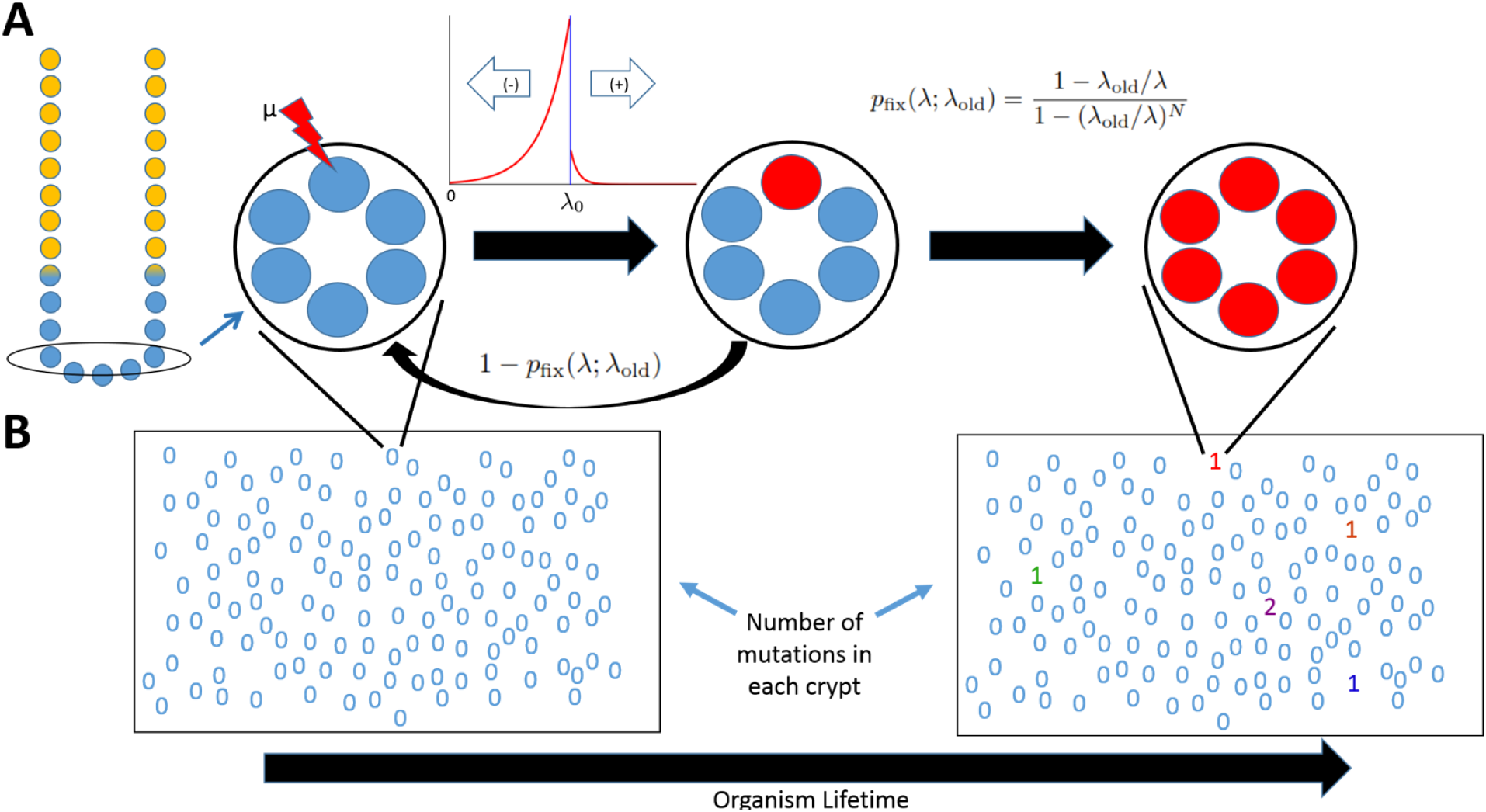
A representation of our model. **A:** A cross section of an intestinal crypt, blue circles at the base of the crypt represent stem cells while yellow circles represent cells that have committed to differentiation. The oval cross section at the base encompasses the stem cell niche, while stem cells above this niche are destined to commit to differentiation. Taking a top-down look at the oval, large circles represent a cross section of the intestinal crypt base, which houses the intestinal stem cells, represented by smaller blue and red circles. Mutations may occur to a single cell in the stem cell niche. These mutations alter the fitness of the cell according to a specified distribution of fitness effects. Given the new fitness, the mutated lineage has a certain probability, *p*_fix_(λ; λ_old_), of reaching fixation within the stem cell niche. **B:** Here, the rectangles represent a cross section of the intestinal epithelium with the numbers representing the locations of individual crypts and describing the number of fixed mutations for each crypt. An organism accumulates fixed mutations over its lifetime.

### Parameter Choices

Some estimates of crypt dynamics parameters have shifted over time, for example the stem cell division rate in the mouse was formerly thought to be once every 1–1.5 days (Lopez-Garcia et al., 2010), but more recent estimates indicate they divide once every 3–10 days (Kozar et al., 2013). Kozar et al. (2013) demonstrated that the division rate of stem cells in the stem cell niche of mice varied from approximately 0.1 to 0.2 to 0.3 divisions per day along the proximal small intestine, distal small intestine, and colon, respectively. Likewise the estimated number of stem cells within the mouse stem cell niche varies from approximately five to six to seven, respectively. The total number of cells in crypts expressing stem cell markers has been reported to be 14–16 in mice (Snippert et al., 2010; Clevers, 2013; Lopez-Garcia et al., 2010). For the analysis of our mouse model we chose the middle value of these parameter ranges, a crypt with 15 total cells expressing stem cell markers, with 6 of the cells constituting the stem cell niche dividing 0.20 times per stem cell per day. To estimate the differentiation rate of stem cells outside the stem cell niche, we used a continuous time Markov chain, described in the Appendix: Description of the mathematical methodology. According to this model, in order for the total stem cells in the crypt of a mouse to stay at a constant population size the differentiation rate of stem cells outside of the stem cell niche must be 0.333 per stem cell per day.

The parameters associated with crypt dynamics in mice have been well described, however, we were unable to obtain any data on population incidence of intestinal polyps or tumors in wild type mice. On the other hand, while crypt dynamics in humans have not been as well-studied, there exists incidence data for large intestine polyps (Chapman, 1963). To parameterize the human colon crypt system we considered a few sources. Nicolas et al. (2007) analyzed the methylation patterns within the human colon crypt and their Bayesian analysis suggests a posterior density mode between 15 and 20 stem cells maintaining homeostasis and constituting the stem cell niche within the crypt. Their distribution is skewed to the left so we chose 20 as an initial value for the number of stem cells within the stem cell niche. Bravo and Axelrod (2013) report an average of 35.7 quiescent stem cells within the human colon crypt through a staining experiment, so we assume there are 36 total stem cells within the human colon crypt. Human colon stem cells divide about once every seven days (Potten et al., 2003), which would mean they would have to differentiate at a rate of about 0.321 per day in order to maintain homeostasis at the assumed initial parameters.

We parameterized the initial DFE based on those measured in whole organisms to evaluate whether they can account for known tumorigenesis incidence. The distribution of fitness effects has been estimated in mutation accumulation experiments and directed mutagenesis experiments. We consider the DFE proposed by Joseph and Hall (2004) in a mutation accumulation study because they report the expected effect size of deleterious and beneficial mutations, as well as the mutation rate and the proportion of mutations that were beneficial in a diploid eukaryotic system (*Saccharomyces cerevisiae*). They report an average beneficial heterozygous fitness effect of 0.061, which is slightly lower but within an order of magnitude of the effect of average beneficial mutation measured for vesicular stomatitis virus of 0.07 (Sanjuán et al., 2004) and *E.coli* of 0.087 (Kassen and Bataillon, 2006). They found that 5.75% of accumulated mutations were beneficial and that the overall mutation rate was 6.3 × 10^−5^ mutations per haploid genome per generation. This would result in a diploid beneficial mutation rate of 2 × 6.3 × 10^−5^ × 0.0575 = 7.245 × 10^−6^. This is within an order of magnitude of the beneficial mutation rate reported for *E. coli* (Wiser et al., 2013).

Mutation accumulation experiments may not capture the true distribution of fitness effects because they rely on observing the mutations of lineages that survive and persist in a population. Because of this, they are biased against mutations of large deleterious effect. Additionally, the random passaging of individuals to repopulate new generations may result in drastically different estimates of average mutational effect size for the same species. For instance, average deleterious effect of mutations in *Saccharomyces cerevisiae* has been estimated to be 0.061 (Joseph and Hall, 2004), 0.086 (Wloch et al., 2001), and 0.217 (Zeyl and DeVisser, 2001). Directed mutagenesis of random genome targets in an RNA virus revealed an average non-lethal deleterious fitness effect of 0.244 (Sanjuán et al., 2004). It is likely that the inherent average effect size of a mutation of deleterious effect would be better reflected by the larger estimates since mutations of large deleterious effect may be lost in mutation accumulation experiments.

After building our model with DFE parameters estimated from whole organisms we describe overall patterns of mutation accumulation and risk of tumorigenesis and then we utilized least squares analysis to explore the best fit among a series of plausible choices of *μ* and the expected value of *s*_+_ for the human incidence curves and compared to data from Chapman (1963) (best fit figures available as Supporting Figures Figure S1, Figure S2, and Figure S3).

The model described above was executed using R version 3.1.1. R scripts developed for this study are available at: to be completed after manuscript is accepted for publication.

**Table 1.** Initial model parameters, combining whole organism DFE with organismal crypt parameters. See text above for reasoning behind initial parameter choices.

| Parameter | Description | Value in mouse<br>[Ref] | Value in human<br>[Ref] |
| --- | --- | --- | --- |
| $P_B$ | Percent of mutations with a beneficial effect | 0.0575 (Joseph and Hall, 2004) | 0.0575 (Joseph and Hall, 2004) |
| $s_+$ | Effect size of a mutation of beneficial effect | 0.061 (Joseph and Hall, 2004) | 0.061 (Joseph and Hall, 2004) |
| $s_-$ | Effect size of a mutation of deleterious effect | 0.217 (Zeyl and DeVisser, 2001) | 0.217 (Zeyl and DeVisser, 2001) |
| $\mu$ | Mutation rate per genes influencing fitness per division | $2 \times 6.3 \times 10^{-5}$ (Joseph and Hall, 2004) | $2 \times 6.3 \times 10^{-5}$ (Joseph and Hall, 2004) |
| $\lambda_0$ | Normal stem cell division rate per day | 0.2 (Kozar et al., 2013) | 0.143 (Potten et al., 2003) |
| $\nu_0$ | Normal stem cell differentiation rate per day | 0.333 (this study) | 0.321 (this study) |
| $N$ | Number of stem cells in the stem cell niche at the base of the crypt | 6 (Kozar et al., 2013) | 20 (Nicolas et al., 2007) |
| $N_T$ | Total number of stem cells expressing stem cell markers in the crypt | 15 (Clevers, 2013) | 36 (Bravo and Axelrod, 2013) |
| Crypts | Number of crypts in the small and large intestine, respectively | $7.5 \times 10^5$, $4.5 \times 10^5$ (Potten et al., 2003) | $5 \times 10^7$, $2 \times 10^7$ (Potten et al., 2003) |

## Results

### Mutations result in both aging and tumorigenesis within the intestine

Because stem cell niche populations are small, it is possible for mutant lineages with a fitness *dis*advantage to fix in the niche. This, coupled with the fact that the vast majority of mutations that occur will have a deleterious effect on stem cell fitness, results in the expected value of the probability density describing the new division rates to move away from the tumorigenesis threshold with subsequent fixed mutations (Fig. 2**A,C,E**). In general, the accumulation of fixed mutations within crypts results in impaired stem cell maintenance and lower stem cell production.

**Figure 2.**
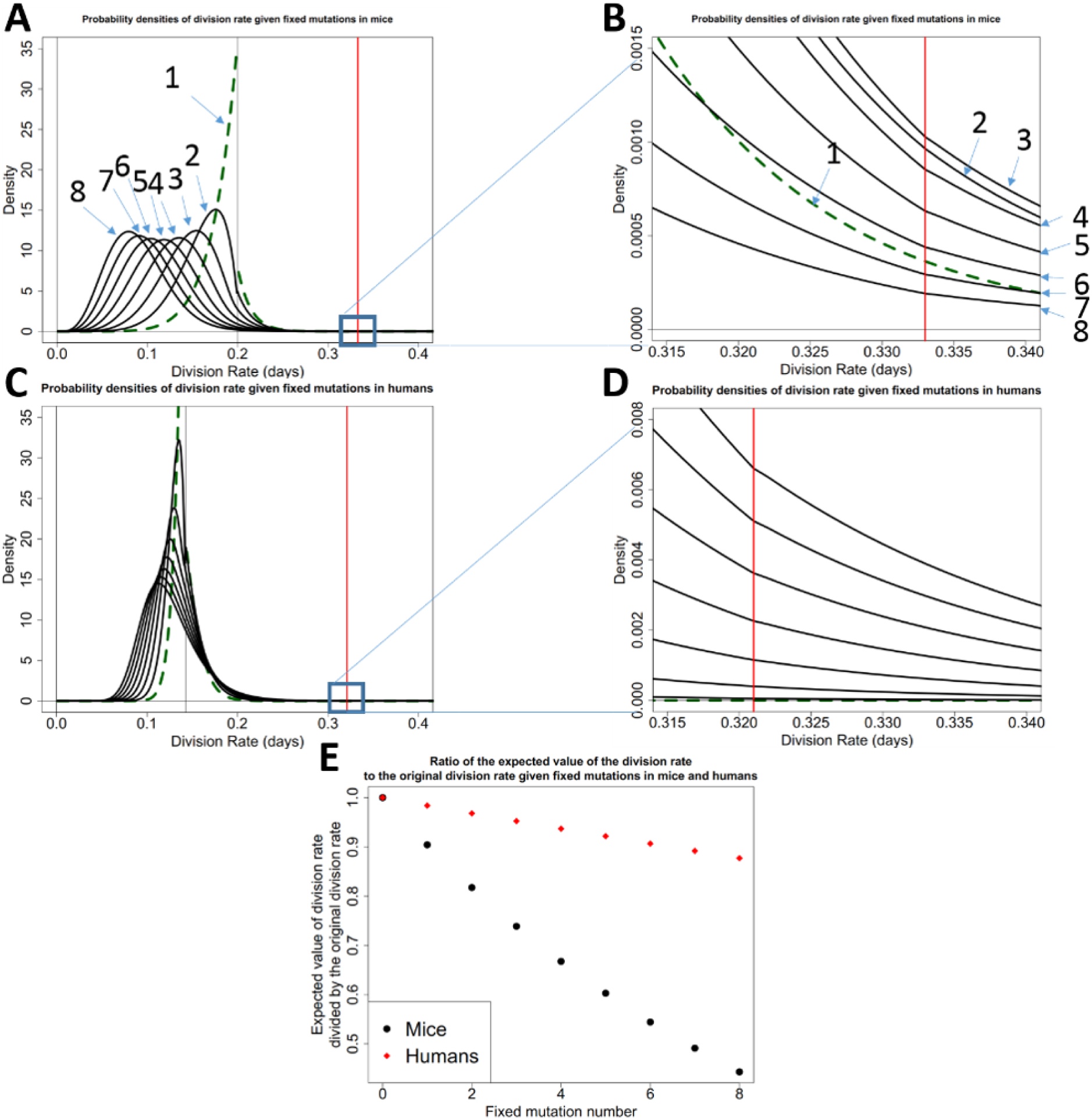
The accumulation of probability densities describing stem cell division rate. **A:** Exponentially distributed fitness effects on division rate using the parameters in Table 1 for the mouse. The first density is a green dashed line. Each probability density represents the division rate of a fixed lineage after *n* fixed mutations, with *n* indicated by an arrow. **B**: Zooming in on the tumorigenesis threshold, we see that the area of the division rate density that is over the tumorigenesis threshold increases at first and then decreases with subsequent mutation. There is a change in slope of the densities at the tumorigenesis threshold because subsequent densities are calculated from the previous density which has had the area to the right of the tumorigenesis threshold removed and the area to the left renormalized to 1. **C,D** are the same as **A** and **B**, respectively, but are for the human scenario. The larger population size decreases the strength of drift. Order of mutations in **C** proceeds as in **A**, and proceeds from 1 through 8 from bottom to top in **D**. **E**: The expected values of the probability densities in **A** and **B** divided by their original values over subsequent fixed mutations

The probability that a particular fixed mutation will result in tumorigenesis in the crypt, *p_n_* (Eq. 5), is equal to the area under these densities that crosses the tumorigenesis threshold (Fig. 2**B,D**). For the initial parameterization in mice and humans this increases at first, but then decreases with subsequent fixed mutations as the probability densities describing division rate move away from the tumorigenesis threshold.

### Predicted incidence curves in mice and humans using DFE derived from a whole organism

Using the model described in “Selection Assumptions”, we determined the cumulative probability distribution of tumorigenesis within a population of crypts in an individual organism. For mice, using the initial parameters in Table 1 and exponentially distributed beneficial fitness effects, we find that the incidence of tumorigenesis is predicted to increase linearly with age, with close to nine percent of mice experiencing tumorigenesis at three years of age (Fig. 3A). Human tumorigenesis incidence in the large intestine is predicted to be approximately 36 percent at 80 years of age (Fig. 3B), using an exponentially distributed beneficial fitness effects and the initial null parameters from Table 1.

**Figure 3.**
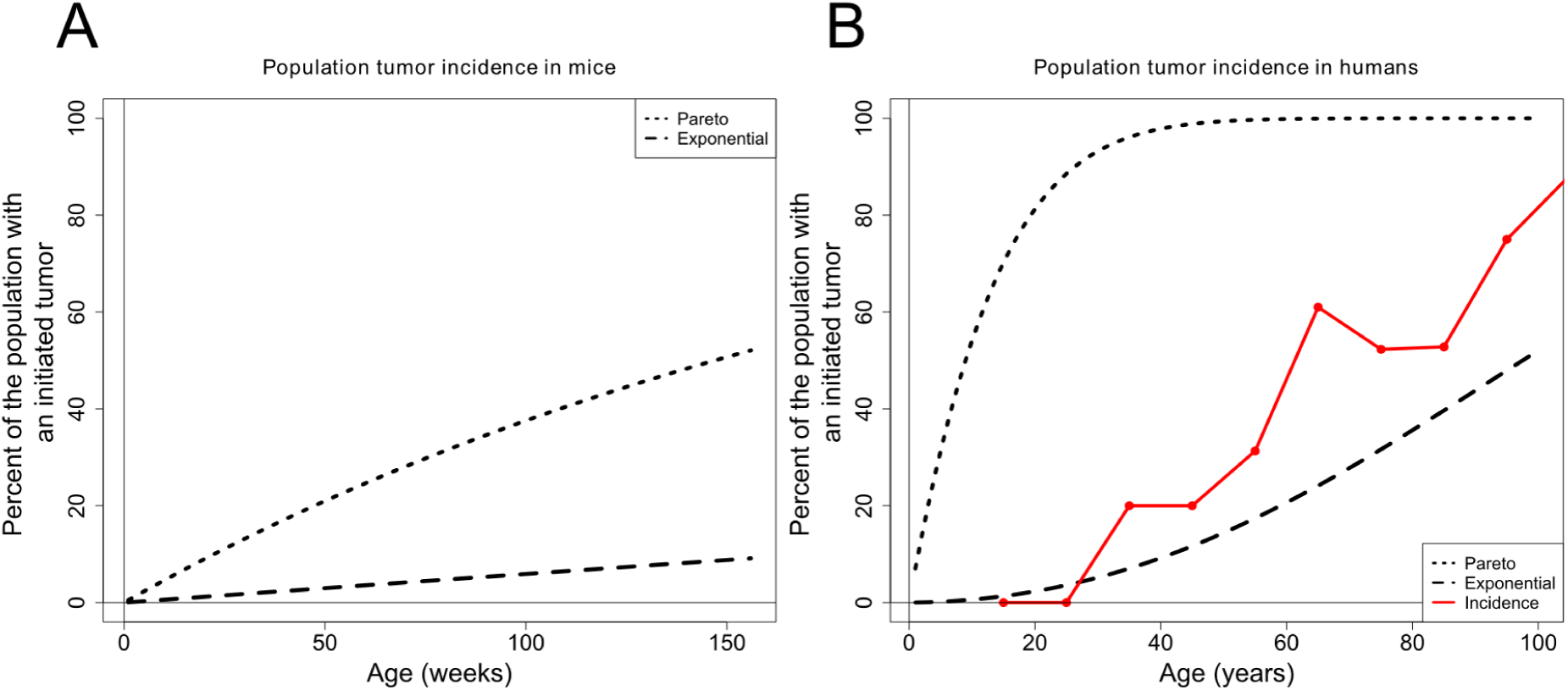
Tumorigenesis incidence in mice and humans using whole organism DFE parameters. **A** The population incidence of tumorigenesis throughout the entire intestinal tract of the mouse. **B:** The population incidence of tumorigenesis throughout the large intestine in humans. The black dashed lines are generated from the species specific parameters listed in Table 1. The solid red line connects large intestine polyp incidence data found during autopsy (Chapman, 1963).

The only incidence data for early tumors or polyps was found for the large intestine in humans. The predicted incidence curve derived from an exponentially distributed DFE follows the same qualitative dynamics as the tumor incidence data. Incidence curves that are derived from a power-law distribution using the initial parameters in Table 1 predict nearly 100 percent tumorigenesis by 80 years of age and do not follow the incidence data dynamics. Hence, we performed a least squares analysis, varying parameters that have not been characterized for human somatic tissue, to find the parameter set in our exploratory space with the best fit to the observed incidence curve to the data.

### Altering the expected beneficial fitness effects and the mutation rate provides better fits for both exponential and power-law derived incidence curves

The expected mean fitness effects (*s_+_,s_−_*) of the DFE and the mutation rate (*μ*) per division of a mutation which alters the stem cell fitness were inferred from whole organisms as an initial parameter choice (Table 1). A parameter space around the initial choices was explored and a least squares analysis was performed to find a better fit to the data (additional information in the Appendix: Description of the mathematical methodology, Supporting Figures Figure S1, Figure S2, and Figure S3). Just the mutation rate (*μ*) and expected beneficial fitness effect (*s*_+_) are presented because changes to the expected deleterious fitness effect (*s*_−_) had little effect on the resultant tumorigenesis incidence curves. We found that both the exponential and power-law scenarios can provide similarly good fits to the data, however with distinctly different parameters. The exponential DFE derived curve provided the best fit with the same mutation rate as our initial choice (Table 1), with a slightly larger expected beneficial fitness effect (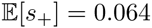, Fig. 4A, red dashed line). Interestingly, assuming the same expected beneficial fitness effect as in Table 1 and varying the mutation rate provides a reasonable fit with a slightly larger mutation rate (*μ* = 1.75 × 10^−4^, 4A, blue dashed line). The power-law DFE provided a similarly good fit to the incidence curve, but for a parameter space that assumes a much smaller expected beneficial fitness effect and a large mutation rate (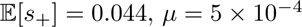, *μ* = 5 × 10^−4^, Fig. 4B red dashed line).

**Figure 4.**
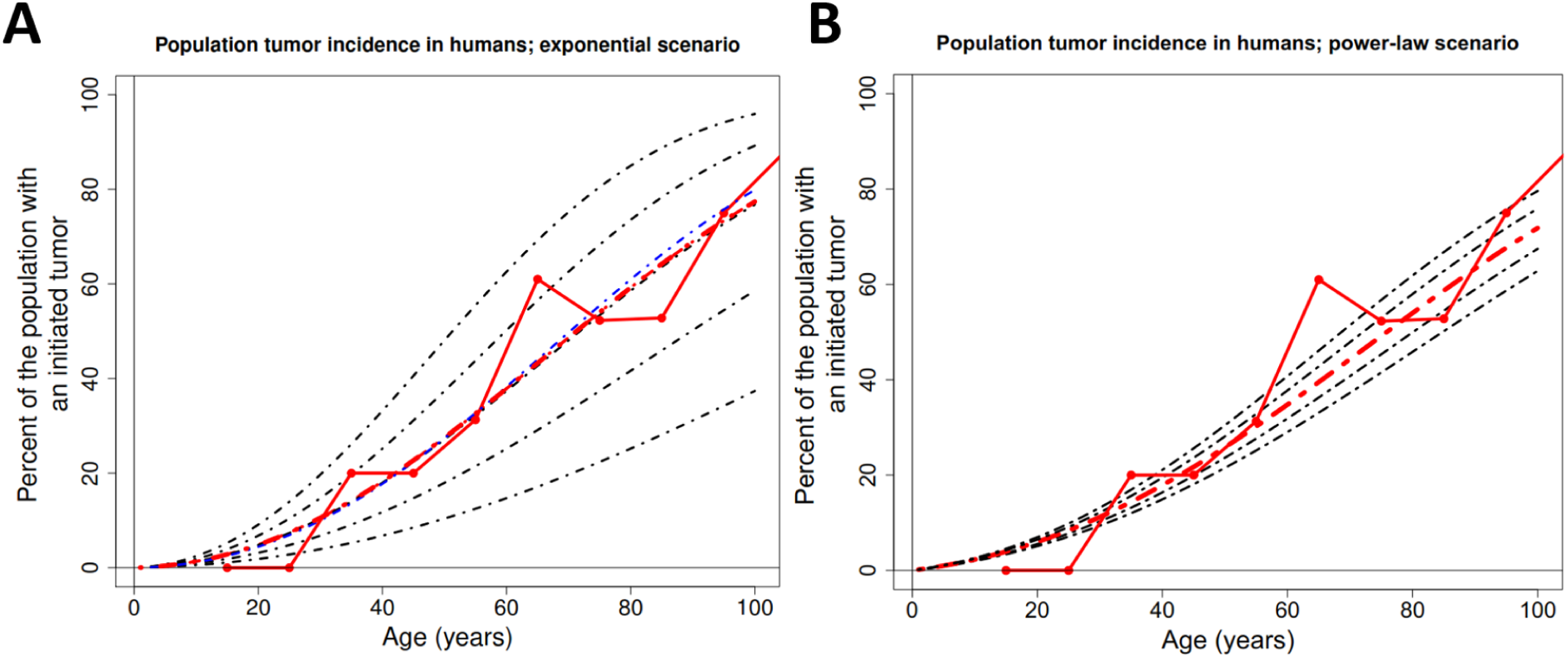
Tumorigenesis incidence curves resulting from least squares parameter fitting. **A:** Incidence curves derived from the assumption of an **exponential beneficial DFE**. The best fit to the data out of the explored parameter space has the same *μ* as the yeast reported in 1 and 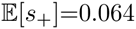 (red dashed line). Black dashed lines derived from 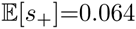 and, from bottom to top, *μ* = 7.5 × 10^−5^ to 1.75 × 10^−4^ by 2.5 × 10^−5^. Blue dashed line is the predicted incidence curve with the best fit with 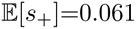 (initial DFE derived from yeast reported in 1), which had *μ* = 1.75 × 10^−4^. **B:** Incidence curves derived from the assumption of a **power-law beneficial DFE**. All parameters are the same as in Table 1, except 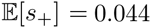 for each curve and, ranging from top to bottom, *μ* ranges from 4.5 × 10^−4^ to 5.5 × 10^−4^ by 2.5 × 10^−5^, with 5 × 10^−4^ providing the best fit.

### Mutations that alter the differentiation rate of stem cells result in rapid aging and tumorigenesis

Mutations affecting differentiation rate influence the lifetime of a stem cell lineage. Mutations that increase differentiation rate will decrease the fitness of the lineage, while mutations that decrease differentiation rate increase fitness. Mutations affecting differentiation rate neutrally drift to fixation in the stem cell niche because the differentiation phenotype is not expressed in the niche, hence all cells divide at the same rate. Thus, the probability of fixation of mutations to differentiation rate is (1*/N*), regardless of mutational effect. We only considered an exponential mutational effect distribution because the distinction between exponential and power law distributions is only significant in prevalence of large deviations from the mean, and beneficial mutational effects in this scenario exist between *ν*_0_ and zero. Because all mutations that solely affect differentiation rate drift neutrally, and the majority of mutations decrease fitness (by increasing differentiation rate), the majority of fixed mutations move stem cell pools away from the tumorigenesis threshold. (Fig. 5).

**Figure 5.**
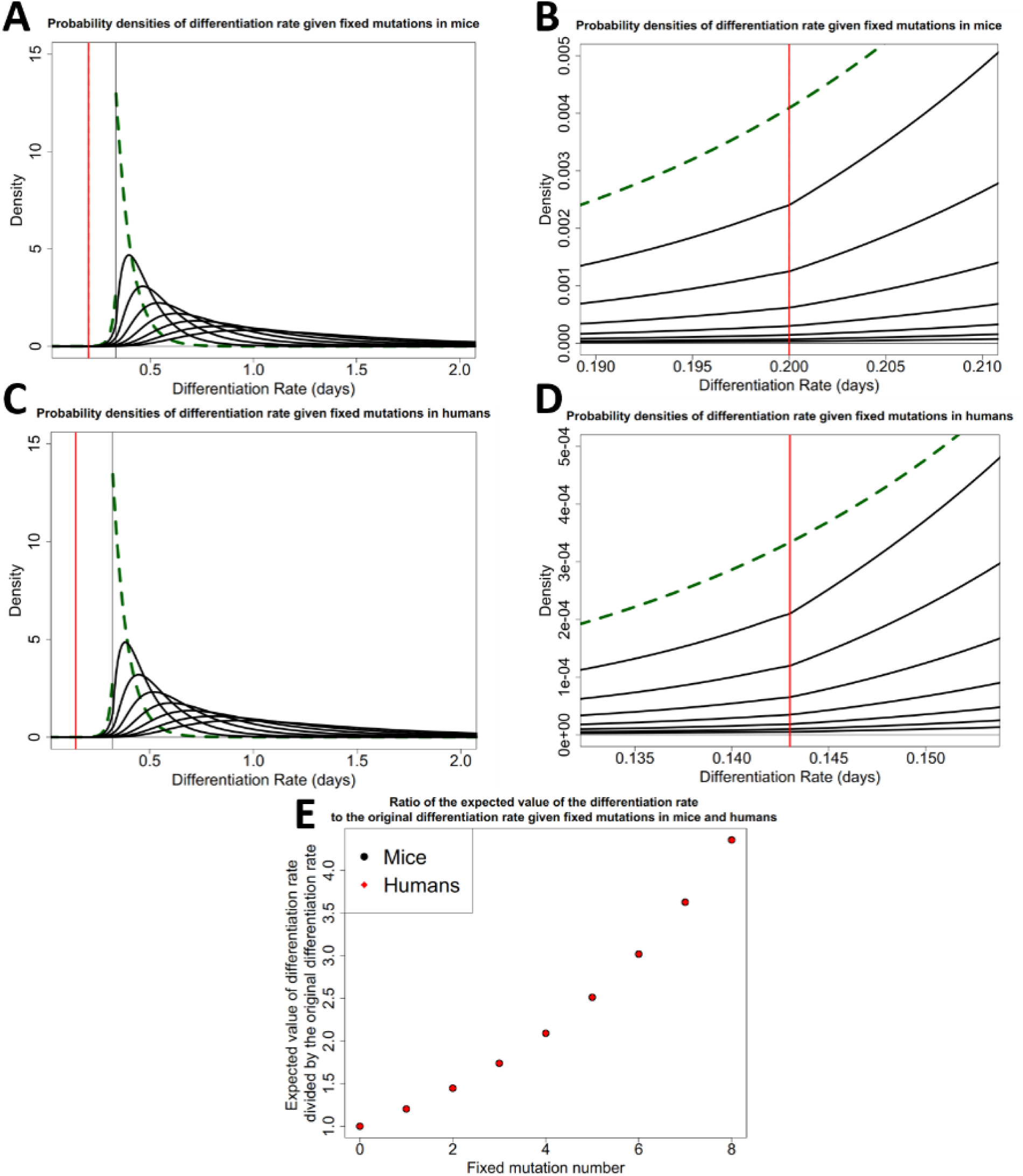
The accumulation of probability densities describing stem cell differentiation rate. **A**: Exponentially distributed fitness effects on differentiation rate using the parameters in Table 1 for the mouse. The first density is a green dashed line. Each probability density represents the differentiation rate of a fixed lineage after *n* fixed mutations, with subsequent mutations traveling away from the original differentiation rate. **B**: Zooming in on the tumorigenesis threshold, we see that the area of the differentiation rate density that is over the tumorigenesis threshold decreases with subsequent mutation. There is a change in slope of the densities at the tumorigenesis threshold because subsequent densities are calculated from the previous density which has had the area to the left of the tumorigenesis threshold removed and the area to the right renormalized to 1. **C,D** are the same as **A** and **B**, respectively, but are for the human scenario. Order of mutations in **C** proceeds as in **A**. **E**: The expected values of the probability densities in **A** and **B** divided by their original values over subsequent fixed mutations

Mutational effects are typically described as a proportion of the phenotype they are affecting, and as such the same DFE applied to a larger rate will have a larger absolute expected effect. The differentiation rate of stem cells displaced from the niche is necessarily larger than the intrinsic division rate since only a subpopulation of the entire stem cell population is exposed to committing to differentiation, however all cells are dividing (Ritsma et al., 2014), and the stem cell population is maintained at a steady-state equilibrium. Thus, mutations affecting differentiation rate in our model have a larger absolute effect for the same proportional change in rate when compared to the previous analysis on mutations to division rate. Hence, given a fixed mutation, we see a high incidence of tumorigenesis when the mutation affects differentiation rate (Fig. 6A,B). Fitting analyses along a range of plausible parameter space revealed a poorer fit when compared to mutations that alter division rate because mutations that alter differentiation rate will always result in large tumor incidence at early age.

**Figure 6.**
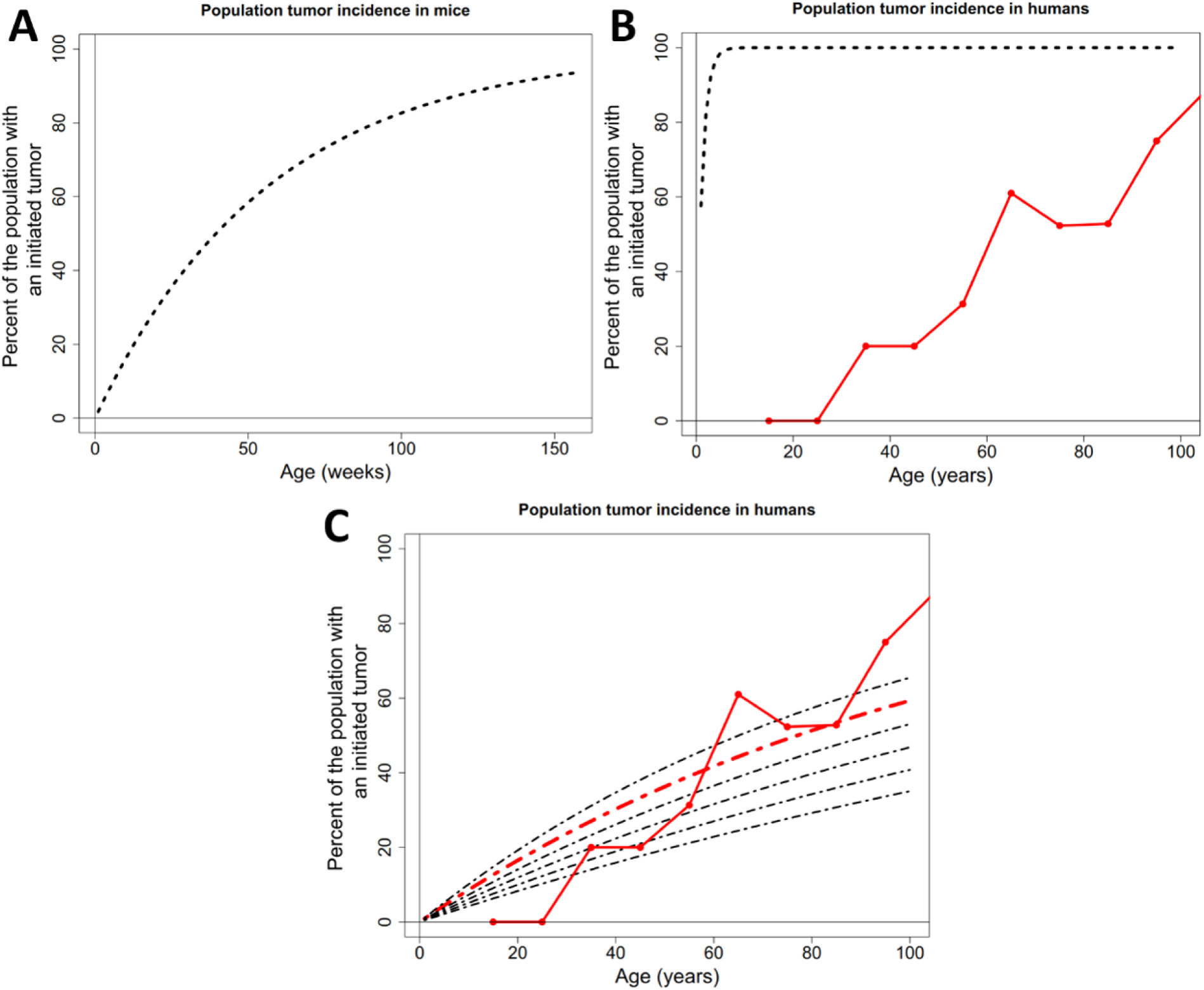
The tumorigenesis incidence resulting from stem cell mutational effects on differentiation rate. Calculations presented for tumor incidence in **A** mice, **B** humans, and **C:** Best fit incidence curve in red; expected beneficial fitness effect of 0.057 and mutation rate of 2.5×10^−6^. The other curves have the same mutation rate but vary around the expected beneficial fitness effect by increments of 0.001.

## Discussion

### Whole organism DFE are sufficient to explain tumorigenesis

We hypothesized that mutations in somatic tissues would differ in their distribution, compared to unicellular whole organisms, because of the regulatory processes that control cell division and differentiation rates in multicellular organisms. However, we found that whole organism DFE were sufficient to account for patterns of tumorigenesis in the intestines. This suggests that somatic evolution is not unique; but instead is based on the same patterns of mutation that we see in whole organisms. Hence, the differences in evolutionary patterns between somatic tissues and whole organisms, such as the tendency of tissues to age via mutation accumulation while populations of whole organisms instead evolve to greater mean fitness in benign environments, arise as a consequence of the small populations of stem cells within multi-cellular organisms and the asexual nature of cell division. Somatic aging via mutation is thus akin to the action of Muller’s ratchet, the accumulation of deleterious mutations in organisms that cannot eliminate them via recombination. Indeed, the ratchet acts more strongly than in populations of organisms as a result of the relative importance of drift versus selection in very small stem cell populations (i.e., niches). This raises the interesting question of why somatic tissues are organized in this way and whether small stem cell pools predominate to minimize tumorigenesis at the expense of aging, as has been suggested by Michor et al. (2003).

Given the role of well-known large effect mutations in cancer, it is tempting, from a mathematical modeling point-of-view, to adopt a heavy-tailed (infinite variance) distribution for the DFE. In contrast to the DFE employed for modeling populations of whole organisms (e.g. an exponential distribution), which tend to exhibit small incremental changes, a heavy-tailed regime enables a significant contribution from “one-shot” large mutations. To probe this possibility, we included in our simulations a power-law (Pareto) distribution which, through its shape parameter *α*, can be either heavy-tailed (1 < *α* ≤ 3) or not (*α >* 3). It is noteworthy, then, that the best fit parameters were very far from the heavy-tailed regime (*α* ≈ 16). The prevalence of large outlier mutations for such a distribution is comparable to what would be seen from an exponential distribution, meaning that the heavy-tailed regime is not an appropriate modeling framework to explain the data.

### Small populations and genetic drift lead to aging

One of our primary findings is that mutation effects drive crypt aging more so than tumorigenesis. Tumor formation and aging are two manifestations of the accumulation of cellular genetic damage. This damage is especially relevant to the aging process when it affects the functional competence of stem cells and compromises their ability to replenish the various cell populations of their constituent tissue (López-Otín et al., 2013). Mutations to stem cells that result in aging have been associated with diminishing the stem cell’s potential to proliferate (Rossi et al., 2007), competitively exclude healthy stem cells (Nijnik et al., 2007), and self-renew or differentiate (Moskalev et al., 2013; Jones and Rando, 2011). These effects on stem cell dynamics would decrease the number of functional stem and non-stem cells in tissues, thus resulting in tissue aging, as defined in aging reviews and experimental work (above) and previous mathematical models (Wodarz, 2007). As mutations become fixed in the intestinal stem cell niche the expected value of the probability density describing new stem cell lineage division rates decreases when we consider mutations that affect division rate, and the expected value for differentiation rate increases when we consider mutations that affect differentiation rate, and thus crypts are predominately aging. The intestinal stem cell niche is maintained at a population size smaller than the effective population sizes of whole organisms and our findings derive directly from this population structure.

Our study, which emphasizes small healthy crypt populations, contrasts with previous studies that have investigated the accumulation of deleterious mutations in somatic tissue. These studies have looked at larger initial population sizes and in effect model hyperplasia or growing tumors. For instance, McFarland et al. (2013) modeled populations with an initial population size of approximately 1000 cells based on estimates from hyperplasia in mice two weeks after *APC* deletion. Similarly, Mcfarland et al. (2014) investigated mutation accumulation in models of hyperplasia and growing cancers. Datta et al. (2013) modeled deleterious mutations in housekeeping genes in an exponentially growing tumor initialized at 1 × 10^6^ cells. Beckman and Loeb (2005) assumed their population of cells was sufficiently large to ensure a deleterious mutation of any strength could not become fixed. These approaches are useful to describe tumor growth in initiated tumors but fail to capture the relative importance of drift in evolving stem cell niches and the process of tumorigenesis from healthy stem cell niches, which exist as very small populations.

We find that crypts with fixed mutations are distributed along a range of both aging and tumor formation. The expected value of the division rate density moving away from the tumorigenesis threshold causes the probability of tumorigenesis per fixed mutation to eventually decrease. For example, in mice, there is a smaller probability that the fourth fixed mutation in a stem cell niche will result in tumorigenesis when compared to the third mutation in the exponential beneficial DFE scenario. The human intestinal crypt stem cell niche consists of a larger number of stem cells so drift plays a smaller role in the evolutionary trajectory of these crypts. Nonetheless, the mode of the distribution of division rate still moves away from the tumorigenesis threshold, albeit at a slower rate. Additionally, the probability of tumorigenesis per fixed mutation reaches a maximum at the twentieth fixed mutation in the case of the human in the exponential beneficial DFE scenario.

Although our model assumes that the size of the stem cell niche (*N*) remains constant and mutations only change the division rate or differentiation rate of lineages, it’s possible that mutations could alter the niche size. If mutations altered the size of a crypt’s stem cell niche they would change the probability of fixation of subsequent mutations.

### Mutations that only affect differentiation rate do not match incidence data curves

Analyses of colon cancer genomes from different individuals reveals that a small number of genes, associated with large fitness advantage, are commonly mutated among cancers (Wood et al., 2007). For instance, many colon cancers contain cells that have mutations in genes involved in the Wnt signaling cascade responsible for maintaining “stemness” (Clevers and Nusse, 2012). A study by Smith et al. (2002) found that 56% of 106 sequenced tumors had mutations in the APC gene, which, when nonfunctional, results in the activation of the Wnt cascade (Reya and Clevers, 2005). Additionally, cancers that have a mutation in the APC gene tend to have the mutation distributed throughout the tumor, suggesting the mutations occurred early in tumor growth (Sottoriva et al., 2015). Because the Wnt signaling cascade is involved with maintaining a stem cell phenotype mutations in this cascade would influence the propensity for stem cells to differentiate. Additionally, Smith *et al*. (2002) also found that 61.3% of colorectal cancers had mutations in *p53*, involved in regulating apoptosis, and 27.4% of colorectal cancers had mutations in *K-ras*, thought to drive cancer growth by accelerating stem cell division and leading to enhanced crypt fission (Snippert et al., 2014).

Our modeling scenario of mutations only having an affect on the differentiation rate of stem cells and having effect sizes equal to those measured in whole organisms results in rapid tumorigenesis, with nearly 100% of human individuals having a polyp in their large intestine at young age. Indeed, individuals with familial adenomatous polyposis (FAP), who already have a germline mutation in one copy of their *APC* gene and only need one mutational hit on the other to form an adenoma, regularly develop adenomas as teenagers (Bozic et al., 2010). Even when we decrease the expected beneficial mutational effect size and decrease the mutation rate in an attempt to better fit the tumorigenesis incidence data, we find that mutations only affecting differentiation rate still result in more tumorigenesis than predicted at young age. However, the data was derived from autopsies on individuals greater than 10 years of age, so data for tumorigenesis is lacking in this age group. Additionally, we modeled scenarios where mutations only affect division or differentiation, nature is certainly more complex and mutation in both differentiation and division rates are likely to co-occur within a crypt population. Indeed, the APC protein discussed above contributes directly or indirectly to cellular division, differentiation, migration, cell orientation, and apoptosis (McCartney and Näthke, 2008; Dikovskaya et al., 2007).

We model stem cell dynamics and mutational effects on those dynamics as a property that is controlled by an individual stem cell’s genome, i.e. a stem cell’s heritable ability to produce or respond to internal signals, or respond to external signals, to divide or differentiate. The external signals regulating stem cell phenotype, such as Wnt signals produced by Paneth cells in the small intestine (Clevers, 2013), are produced by cells differentiated from stem cells. Mutations to the stem cell genome may eventually influence the production of these signals in daughter cells. These mutations would drift neutrally in the niche, as they are not expressed until after stem cell differentiation, and, unless the lineage harboring the mutation reaches fixation in the niche, would eventually be lost from the crypt since Paneth cells die in approximately 20 days (Bry et al., 1994). Thus, mutations that result in differential signaling output by daughter cells can be modeled as neutrally fixed mutations acting intrinsically in the stem cells.

### The influence of organism specific factors on somatic evolution

We find less tumor incidence in mice than humans throughout their respective lifetimes using the same DFE parameters. Mice only live a few years and have an order of magnitude fewer crypts in their entire intestine than humans have in just their large intestine (Potten et al., 2003). They also have smaller numbers of stem cells within their crypts, although those stem cells are dividing at a faster rate than human stem cells. Overall, this results in a lower chance of mutant lineages reaching fixation within crypts during the shorter mouse lifetime, and therefore a reduction in the overall number of crypts with fixed mutations is lower. For instance, using the distribution of fixed mutations derived in the Appendix: Description of the mathematical methodology, at two years old a mouse is expected to have about 75 crypts with two mutations, and only about 28 percent of mice will have a single crypt with three mutations. At 85 years old, a human is expected to have about 44 crypts with five mutations, one crypt with six mutations, and about four percent of humans at 85 years old will have a crypt with seven fixed mutations. As humans age they experience more fixed mutations, each of which confers a higher probability of tumorigenesis than the previous, whereas mice are expected to experience the accumulation of fewer mutations, possibly explaining the near linearity of the mouse incidence curve and the upwards curvature of the human incidence curve. Of note, given that a tumorigenesis event has occurred, it is likely the product of one mutation in the mouse model, whereas multiple mutations may contribute to the initiation of a tumor in the human model (the Appendix: Description of the mathematical methodology, Figure S4).

The incidence of polyps at autopsy reported by Chapman (1963) was based on visual observations of discernible elevations of the mucosa in the entire large intestine during autopsy. It would take time for an initiated tumor to grow to a visible mass, so the true tumorigenesis incidence curve may lie in front of the data recorded in this study, with a lag time of growth before the tumor is visible. This lag time would be a function of the individual mutational spectrum of the initiated tumor and the tumor’s environment.

Overall, we have shown that small homeostatic populations of stem cells, typical of somatic tissues in multicellular organisms, accumulate mutations that affect cellular fitness, contributing both to aging and tumorigenesis over an organism’s lifetime. We show that the evolution of intestinal stem cell populations under the assumption of an organismal DFE, as opposed to the assumption of a heavy-tailed beneficial DFE, best predicted early tumor formation. However, aging, rather than tumorigenesis, predominated among crypts in the intestine. Our modeling approach emphasizes tumorigenesis in the context of aging, and vice versa, and demonstrates the importance of mutational processes within very small populations in both these phenomena.

## Acknowledgments

We thank Charlie Baer, Andres J. Garcia, and Jake M. Ferguson for useful discussion. This research was partially supported by the National Science Foundation under Grant No. 0801544 in the Quantitative Spatial Ecology, Evolution and Environment Program at the University of Florida.

## Appendix: Description of the mathematical methodology

In this section we describe the mathematical model underlying our research approach. It is a multi-scale model, including the dynamics of the stem cell niche; the consequences for the larger stem cell population and a crypt; the dynamics in a population of crypts that comprise an individual’s colon; and the dynamics of tumorigenesis among many individuals in a population. Due to the very large number of crypts in the colon and the desire to analyze population level incidence curves, we used several principles of rare-event analysis in our numerical computations and also introduced a few approximations to make computations tractable.

**Population dynamics within the stem cell niche** First, we develop a model for the population dynamics of a crypt immediately after a mutation has occurred. Suppose that there are *N* cells in the stem cell niche and let *X*(*t*) represent the number of cells that are descended from the original mutated cell at time *t*. We model *X*(*t*) as a continuous time Markov chain (CTMC) that takes its values in the set {0,…, *N*} with *X*(0) = 1.

In accordance with the Markov process assumption, the time between divisions of a given stem cell are independent of all other cells and exponentially distributed with rate parameters λ_old_ or λ for the old and the new lineages, respectively. When a cell divides, we assume that there is crowding in the stem cell niche and an old cell is forced out. (In fact, the actual order of events remains unclear. It has also been hypothesized that a cell may leave the niche first, then triggering a cell division to replace it (Lopez-Garcia et al., 2010).) Whether or not the value of the process *X*(*t*) changes depends on whether the cell that has been forced out is from the same lineage as the one that divided. The assumption that leads to the simplest mathematical model is *nearest neighbor displacement*. There are two cases: 1) the dividing cell is of the same lineage as both of its neighbors; and 2) the dividing cell is adjacent to a cell of the opposing lineage. In the first case, the value of *X*(*t*) does not change as a result of the cell division. In the latter case, there is a one-half probability that a cell of the opposing lineage will be displaced. As such, for *X*(*t*) ∈ {1, 2,…, *N* − 1}, the Markov transition rates are given by

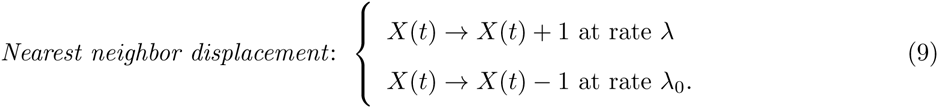

(The rate of one of the two border cells dividing is 2λ for the new lineage and 2λ_old_ for the old lineage and then each is multiplied by the one-half probability of displacing an opposing lineage cell.) An alternate hypothesis is that after division, *any* other cell in the crypt might be displaced. The corresponding transition rates would be

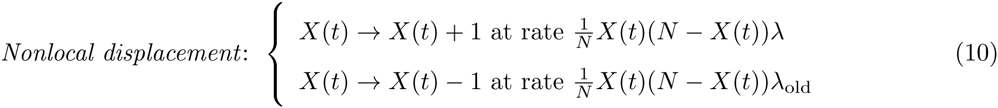

The probability of fixation is actually the same for both models (though the expected time until fixation will differ). Let {*t*_1_*, t*_2_,…} be the sequence of times when *X*(*t*) changes values. Disregarding the role of time in the process, we track the values with the process {*X_n_*}*_n_*_≥0_ defined by *X_n_*:= *X*(*t_n_*). The probability of a transition *X* → *X* + 1 is the rate at which the size of the mutant lineage increases divided by the total rate of change in lineage count. For both models this probability of an increase in the mutant lineage size is *p* = λ/(λ + λ_old_). Using the classical theory of hitting probabilities for biased random walks (Wodarz and Komarova, 2005), one can readily derive the probability *p*_fix_(λ; λ_old_) recorded in Eq. 3 in the main text.

**The intervals between mutations that fix in the stem cell niche**. The DFEs used in this work are both considered in terms of percentage increase or decrease, rather than in terms of absolute quantities of change. In mathematical terms, this means that the densities can be expressed in terms of the ratio λ/λ_old_. A remarkable consequence of this assumption is that the probability of a new lineage fixing in the niche is *independent* of the prevailing division rate λ_old_. To see this, consider the probability of that a new lineage fixes after a mutation drawn from the exponential DFE. Recalling Equation 3, we note that the probability of fixation formula can be written in terms of the ratio of the new to the old division rate, *r* = λ/λ_old_,

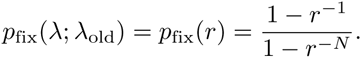

We can then write

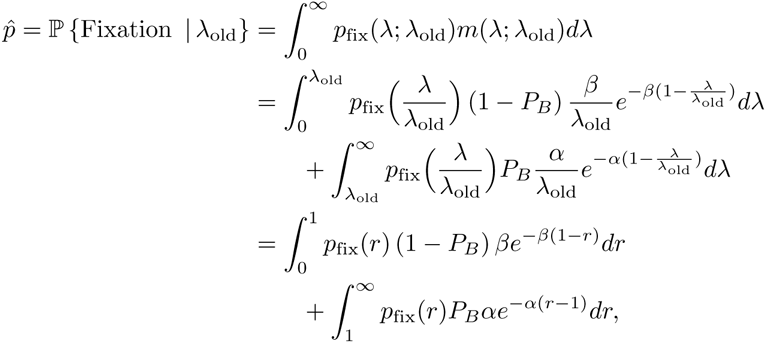
which is independent of the choice of value λ_old_. A similar result holds for the power law DFE. Generally, this property holds for any DFE that can be expressed in terms of the ratio λ/λ_old_. It follows that the number of mutations that must occur in order for a new division rate to fix is distributed Geometrically with success probability 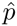. By standard properties of CTMC, we can then say that the time between the arrivals of “successful” mutations is Exponentially distributed with rate parameter 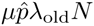.

**Population dynamics outside the stem cell niche**. Once outside the niche, a stem cell can either divide (at rate λ), or it can differentiate into transient amplifying cells (at rate *ν*). For the purposes of this model, we consider differentiated cells to be dead. There is a chance that the lineage of a stem cell outside the niche can undergo sufficiently many mutations to cause tumorigenesis, but we found by way of numerical investigations that this does not significantly contribute to overall incidence of these cancers. As such, let *Y*(*t*) denote the number of stem cells outside the niche that have not yet differentiated. Assuming, for the moment, that all members of the stem cell niche have a division rate λ, the CMTC *Y*(*t*) is defined by the transition rates

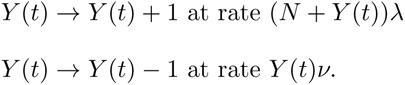

The form of the rate of increase follows from the observation that *Y*(*t*) increases anytime a stem cell divides, whether that stem cell is in the crypt or not. On the other hand, since stem cells in the niche are assumed to not differentiate, the total rate of decrease is proportional to the number of stem cells outside the niche. Because the population size is so small, there is high variability and we note that *Y*(*t*) can regularly hit the value zero. Because the niche is protected by unrelated biological processes, this does not constitute extinction of the full stem cell population. As soon as another stem cell in the niche divides, the population outside the niche is renewed again. A typical trace for *Y*(*t*) can be seen in Supporting Information Figure S5.

The law of this CTMC, *y_n_*(*t*) = ℙ {*Y*(*t*) = *n*}, satisfies the system of master equations

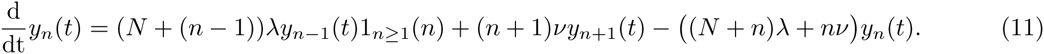

One can then show that the mean 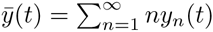 satisfies the ODE

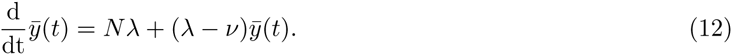

If λ < *ν*, this ODE converges to a steady-state value *Nλ/*(*v −* λ). Otherwise the mean diverges to infinity with exponential growth. For this reason, we consider this threshold to be the initiation of tumorigenesis.

An alternate way to view the dynamics is to note that each time a stem cell in the niche divides it creates a new independent lineage outside the crypt. Let *Y^j^*(*t*) be the number of living stem cells outside the crypt that are descended from (and include) the product of the *j*th stem cell division in the niche. As such 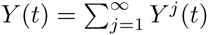. Each process *Y^j^*(*t*) can be understood as a branching process, with an offspring distribution that is Geometrically distributed with “success probability” *q = ν/*(*ν + λ*). (The number of offspring is determined by the number of times the cell divides before differentiating. This is a sequence of independent trials where the probability of having another offspring, rather than differentiating, is λ/(*ν* + λ).) As long as the mean of this offspring distribution is less than or equal to one, these lineages will eventually go extinct. Therefore, the critical stem cell division rate corresponds to when the mean of the offspring distribution (which can be shown to be (1 − *q*)/*q* = λ/*ν*) is less than one. In other words, the critical division rate λ_*_ is simply λ_*_ = *ν*.

**Population dynamics in the crypt**. Of course, tumorigenesis in a given crypt is exceedingly unlikely, even over the lifetime of an individual. We model the colon as a collection of *C* ≈ 10^7^ individual crypts that are mathematically identical and independent. The number of *fixed mutations* in the *i*th crypt at time *t* is denoted *M^i^*(*t*), and let 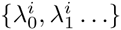 denote the sequence of division rates that become fixed in the *i*th crypt at times 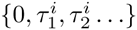 respectively. It follows that the inter-arrival times are independent and distributed as 

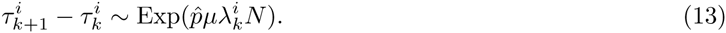

Whether tumorigenesis has occurred in the *i*th crypt will be tracked by the function χ*^i^*(*t*), defined by 

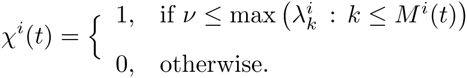

That is to say, χ*^i^*(*t*) = 1 if tumorigenesis has occurred before time *t*. It follows that the time of first tumorigenesis in the colon is given by the time 

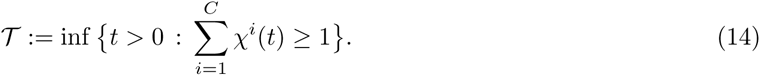

The per capita population incidence curves are then just the cumulative distribution function of the random variable 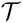, which can be expressed in terms of the individual crypt dynamics as follows:

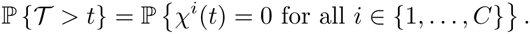

To prepare for our numerical approximation of this quantity we introduce one last bit of notation, 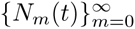, which represents the number of crypts that have seen the arrival of *m* new fixed lineages as of time *t*. Then

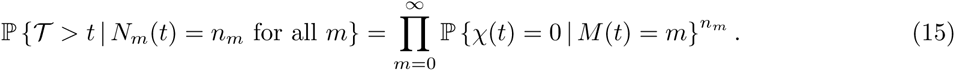

These dynamics can be simulated by Gillespie’s method (Gillespie, 1977), but such an approach is computationally intensive. For this reason we introduced a few simplifying assumptions. For example, we model the arrival rates of new fixed lineages in the crypts as being constant over time (having fixed rate 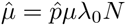, rather than a sequence of rates given in Eq. (13)). This allows us to assume that the number of mutations in each crypt at time *t* is Poisson distributed with mean 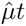. With such a tremendously large number of crypts in the colon, it is in turn reasonable to assume that the number of crypts takes the form 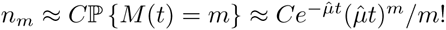. To complete the derivation of Eq. 8 in the main text, we truncate the infinite product in Eq. (15) and note that in the notation of the main text,ℙ {χ(*t*) = 0 | *M*(*t*) = *m*} = *q_m_*.

DFE equations. To define the parameters of the system, we specified the probability *P_B_* of a beneficial (versus deleterious) mutation and the respective means *s*_+_ and *s*_−_ of the DFE *conditioned* on the event that the mutation is beneficial or deleterious. The form of the mean of the DFE depends on whether it is exponential or heavy-tailed. The exponential form DFE has mean

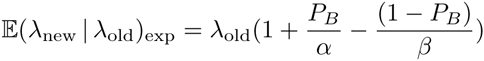
while the heavy-tail DFE has mean

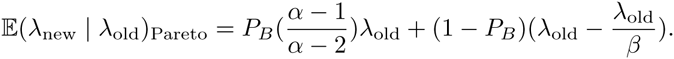

The conditional means have the form

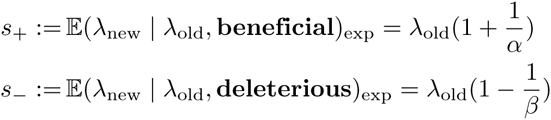
for the exponential DFE, and

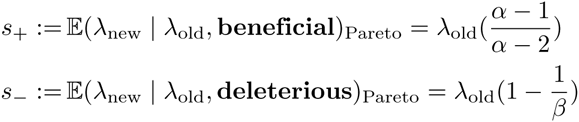
for the heavy-tail DFE.

Least squares analysis. We generated tumor incidence curves and used a least squares analysis to determine which set of these parameters best fit the tumor incidence data described in Chapman (1963). The best fit has the smallest sum of squared residuals of the parameter space explored in figures S1, S2, and S3.

Tumor mutational profile Supporting Information Figure S4 contains the probabilities that each individual mutational profile was the culprit in tumorigenesis given that tumorigenesis occurred in a single crypt. They were calculated by using Bayes’ Theorem to compute the probability that a certain mutational load fixed in the crypt given that a tumorigenesis event happened,

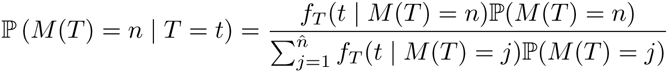
where we recall that *M*(*t*) is the number of fixed mutations as of time *t*, *T* is the precise time that tumorigenesis occurs in the crypt and *f_T_* refers to density of the random variable *T*. The quantity ℙ(*M*(*T*) = *n*) is the probability that tumorigenesis occurs exactly on the nth mutation, a quantity we defined earlier as *p_n_* and gave a recursive formula for in Equation 5 in the main text. To compute the quantity *f_T_*(*t* | *M*(*T*) = *n*), note that since the arrival time of the nth mutation is independent of the event that it causes tumorigenesis, we have that 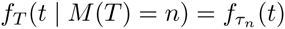, where *τ_n_* is the arrival time of the nth mutation. By hypothesis, *τ_n_* is Poisson distributed with mean 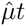 with 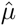 being defined after Equation 8 in the main text. The distributions of mutational profiles given the tumorigenesis event occurred at a certain point in time throughout an organism’s lifetime are given in Figure S4.

**Figure S1.**
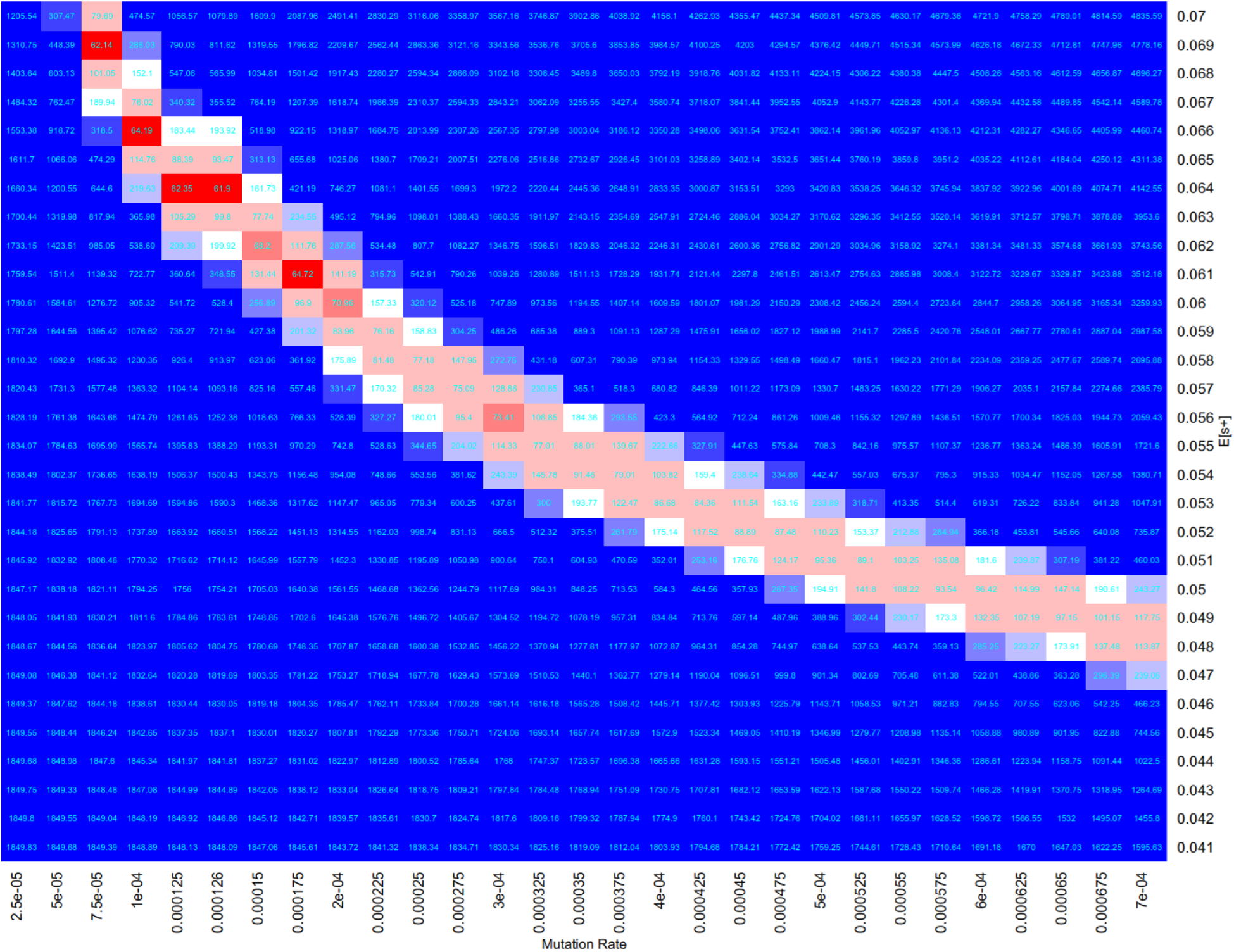
Heat map depicting values for the least squares analysis of predicted tumor incidence and human tumor incidence data for the exponential beneficial DFE on division rate scenario. Parameter combinations with red colors have the smallest sum of squared residuals, while blue colors have the largest.

**Figure S2.**
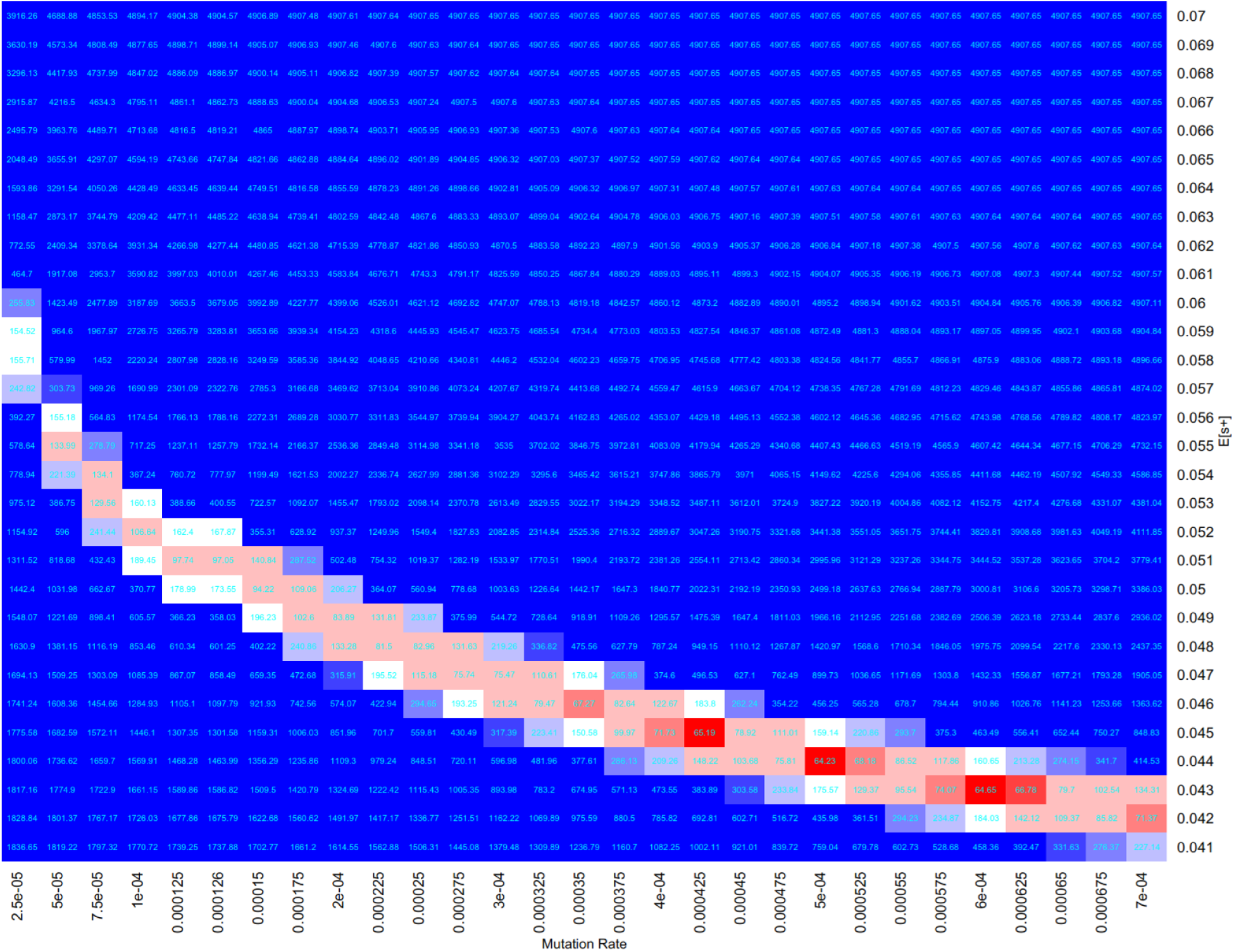
Heat map depicting values for the least squares analysis of predicted tumor incidence and human tumor incidence data for the power-law beneficial DFE on division rate scenario. Parameter combinations with red colors have the smallest sum of squared residuals, while blue colors have the largest.

**Figure S3.**
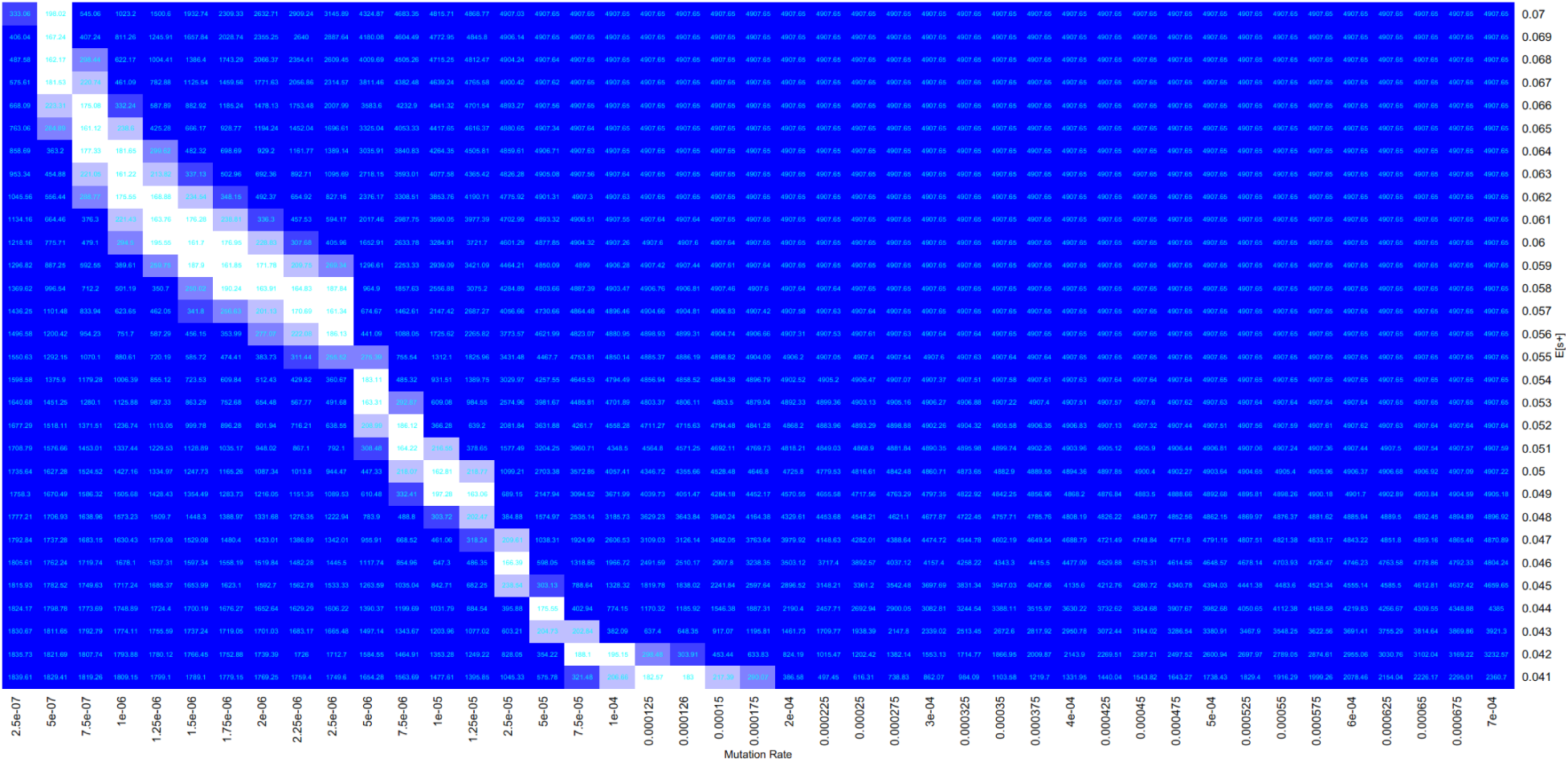
Heat map depicting values for the least squares analysis of predicted tumor incidence and human tumor incidence data for the mutations affecting differentiation rate scenario. Parameter combinations with white colors have the smallest sum of squared residuals, while blue colors have the largest.

**Figure S4.**
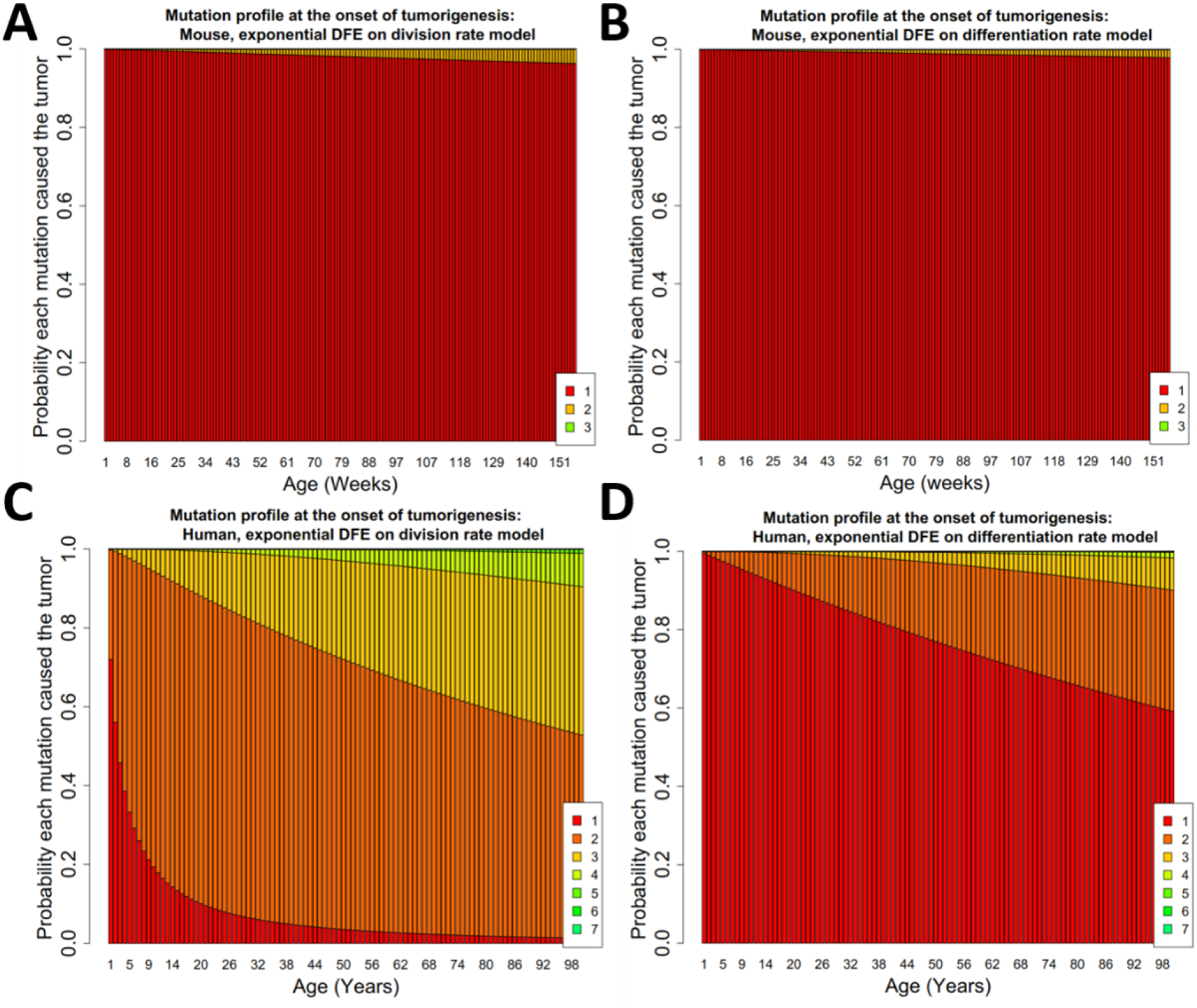
Mutation profiles of a tumor at the onset of tumorigenesis. Probabilities describing the distributions of mutations that caused an initiated tumor throughout the lifetime of a mouse (**A,B**) and human (**C,D**) under the mutations affecting division rate (**A,C**) and differentiation rate (**B,D**) modeling scenarios.

**Figure S5.**
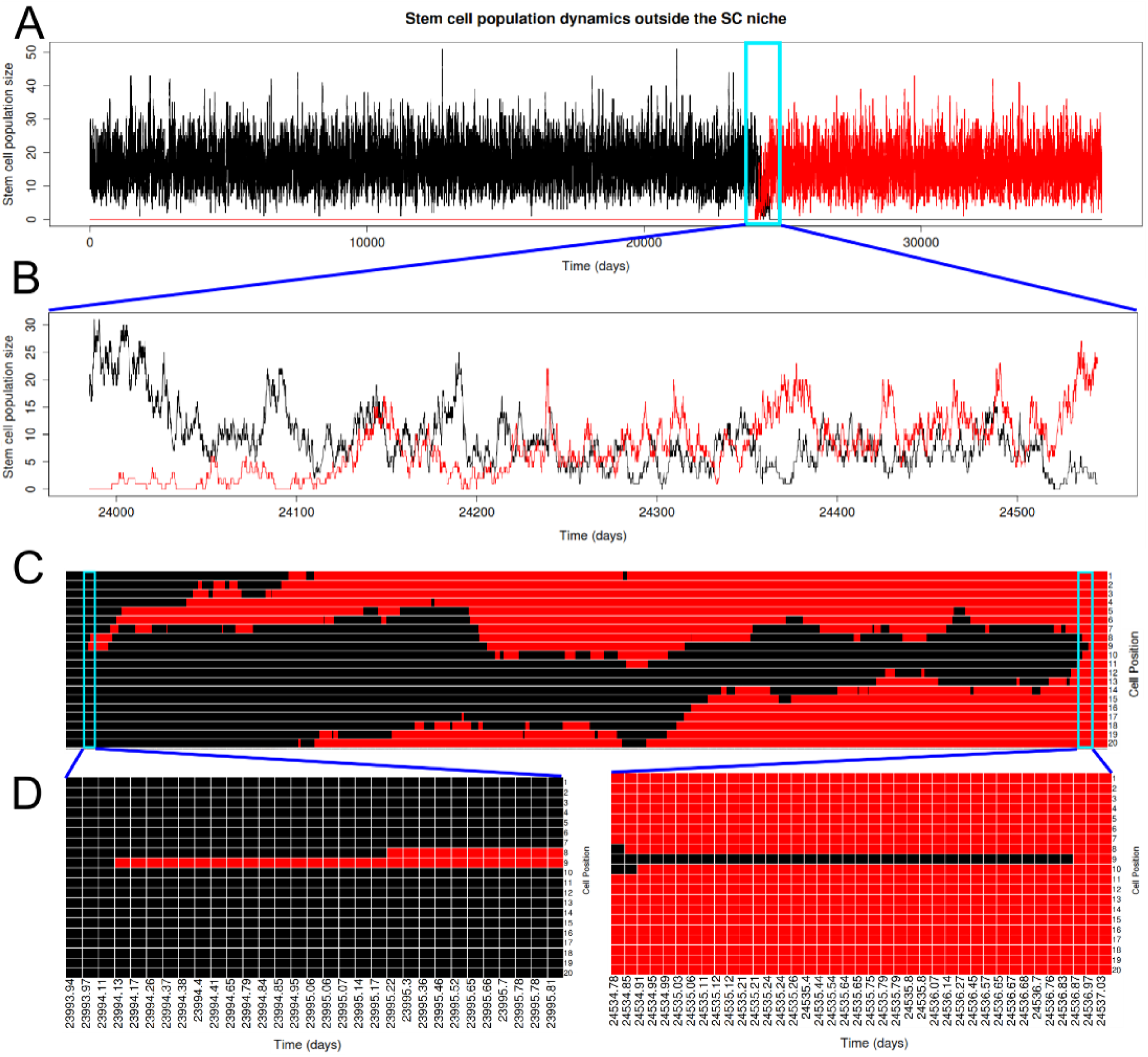
The simulated stem cell dynamics within a human crypt for both the displaced stem cells (A,B) and the stem cell niche (C,D) showing a fixation event of a mutant lineage. Original division rate λ_0_=0.143, new fixed division rate λ_1_ =0.141. **A:** The displaced stem cell population size fluctuates stochastically as stem cells enter the population by being displaced from the niche and leave by committing to differentiation. **B:** Zooming in at the moment a new stem cell lineage begins being displaced from the niche, we see the original (represented in black) lineage going extinct as the new (represented in red) lineage eventually dominates the population. **C:** The spatially explicit stem cell niche is arranged in a circle with cells displacing their neighbors through division. Here that circle is opened (cell positions 1–20 represented on the Y-axis and shown through the same time series as **B**. Each time point corresponds to any event occurring in the entire simulation (division in the stem cell niche, division and differentiation occurring in displaced cells). A new mutant lineage arises through errors caused by division at time 23994.13 days and, by stochastically dividing and displacing neighbors, reaches fixation in the population by day 24536.87. The beginning and end dynamics of this new lineage are shown in inset D

